# Functional triplet motifs underlie accurate predictions of single-trial responses in populations of tuned and untuned V1 neurons

**DOI:** 10.1101/238345

**Authors:** Joseph B Dechery, Jason N MacLean

## Abstract

Visual stimuli are encoded in the activity patterns of neocortical neuronal populations. Trial-averaged neuronal activity is selectively modulated by particular visual stimulus parameters, such as the direction of a moving bar of light, resulting in well-defined tuning properties. However, a large number of neurons in visual cortex remain unmodulated by any given stimulus parameter, and the role of this untuned population is not well understood. Here, we use two-photon calcium imaging to record, in an unbiased manner, from large populations of layer 2/3 excitatory neurons in mouse primary visual cortex to describe co-varying activity on single trials in populations consisting of tuned and untuned neurons. Specifically, we summarize pairwise covariability with an asymmetric partial correlation coefficient, allowing us to analyze the population correlation structure with graph theory. Using the graph neighbors of a neuron, we find that the local population, including tuned and untuned neurons, are able to predict individual neuron activity on a single trial basis and recapitulate average tuning properties of tuned neurons. We also find that a specific functional triplet motif in the graph results in the best predictions, suggesting a signature of informative correlations in these populations. Variance explained in total population activity scales with the number of neurons imaged, suggesting larger sample sizes are required to fully capture local network interactions. In summary, we show that unbiased sampling of the local population can explain single trial response variability as well as trial-averaged tuning properties in V1, and the ability to predict responses is tied to the occurrence of a functional triplet motif.

**Author summary:** V1 populations have historically been characterized by single cell response properties and pairwise co-variability. Many cells, however, do not show obvious dependencies to a given stimulus or behavioral task, and have consequently gone unanalyzed. We densely record from V1 populations to measure how trial-to-trial response variability relates to these previously understudied neurons. We find that individual neurons, regardless of response properties, are inextricably dependent on the population in which they are embedded. By studying patterns of correlations between groups of neurons, we identify a specific triplet motif that predicts this dependence on local population activity. Only by studying large populations simultaneously were we able to find an emergent property of this information. These results imply that understanding how the visual system operates with substantial trial-to-trial variability will necessitate a network perspective that accounts for both visual stimuli and activity in the local population.

## Introduction

In the visual system, decades of research have unraveled the stimulus parameters underlying trial-averaged response properties in single neurons [1]. These response properties have revealed principles of functional organization in primary visual cortex (V1), such as orientation columns [2], and canonical computations, such as divisive normalization [3]. However, responses are variable across trials [4], making the relationship between relatively stable perception and single-trial stimulus representations unclear [5]. The fluctuations of response strength are not independent across neurons, and this shared variability impacts population representations of visual stimuli [6–7]. Neurons are highly interconnected and connection likelihood is biased toward spatially proximal neurons [8], suggesting that trial-to-trial response variability may be in part the manifestation of the state of the surrounding neuronal population [9–10]. Pairwise interactions within a population can shape information representation [11–13], and can be regulated by top-down influences [14]. Therefore, comprehensive descriptions of stimulus representations in primary sensory cortex require a network perspective. Here, we used two-photon imaging to record from large populations of L2/3 excitatory neurons in mouse V1 to study effects of local population activity on trial-to-trial variability.

Understanding the sources and consequences of response variability is necessary to extend theories of sensory computation from the average case to single trials. Perception and behavior take place in real time, after all, so variable responses must be taken into account to understand stimulus representations in cortex. Shared variability in neural responses is commonly quantified by the set of pairwise correlations between neurons, and the structure of these correlations can have constructive or destructive effects on stimulus encoding in populations of neurons [15–16], highlighting the importance of its characterization. Moreover, complex patterns of population activity in retina can be captured by taking into account only neuron firing rates and pairwise correlations [17]. Covariability can also be shaped by cognitive properties such as attention in order to improve perceptual acuity [14]. Whether trial-to-trial variability is harnessed to improve the fidelity of sensory representations, or accounted for when decoding from noisy signals, the properties of response variability have a large impact on neural function.

Research on the correlation structure of population activity is still incomplete, however, and can be expanded by incorporating a more comprehensive sampling of the network [18]. V1 populations consist of neurons whose activity is not modulated by, or is untuned to, a given stimulus. It is still an open question how this subpopulation contributes to neuron correlations. Two-photon imaging results in a relatively unbiased sampling of spatially proximal neurons including both tuned and untuned subpopulations. Neurons unrelated to a behavioral task can help predict activity in neighboring neurons in hippocampal CA1 [9]. In V1, it has been shown that untuned neurons can help to decode the orientation of drifting gratings [19]. We investigate how cofluctuations in the activity within tuned and untuned neurons interacts with responses to drifting grating stimuli.

We characterize population activity and correlations between tuned and untuned subpopulations in order to understand the relationship between single-cell response properties and recurrent network dynamics. Traditional noise correlation analyses study covariability independent from stimulus-driven activity [20]. However, untuned neurons have no stimulus modulation, so we use an analogous partial-correlation based method that additionally accounts for population-wide covariability. Fluctuations common across a local population are an important determinant of single-trial responses in mouse V1 [21–22]. Capturing this additional variable allows us to study the correlations in the entire unbiased sampling of the population. Additionally, the partial correlation matrices are asymmetric and relatively sparse, and can thus be represented as a weighted, directed graph. Graph theory analysis is used to summarize structure in complex networks, and can resolve emergent properties resulting from pairwise relationships. Connectivity patterns, or motifs, in graphs can be characterized within small groups of neurons [23] or across an entire population [24–25]. Motifs have proved to be impactful for understanding complex biological processes including transcription networks [26] and spike propagation [23]. More generally, motifs patterns impact information representation in complex systems [27–28] and have increasingly been a subject of interest among neuroscientific disciplines.

From the graph neighbors of a given neuron, we can accurately predict activity on single trials using a simple, linear model. The local population contains information sufficient to predict trial-to-trial variability and recapitulates average tuning properties. Furthermore, neurons that are well-modeled by the activity of their neighbors have specific signatures of functional connection motifs. Across the entire graph, the most predictive motifs are also the most prevalent, suggesting that this structure is responsible for the overall quality of reconstruction observed. The triplet motif that facilitates predictions of neural activity may have a broad impact on information representation in graphs. Notably, total variance explained in a field of view scales with the number of neurons imaged, suggesting larger sample sizes are required to fully capture local network interactions.

## Results

### Response properties of V1 populations

To study interactions and response variability in local, cortical populations (<800um diameter imaging plane), we imaged L2/3 excitatory neurons (72–347 neurons; 25–33 Hz; n=8 animals; 23 distinct fields of view; Fig 1A) in mouse V1 during presentation of drifting gratings (Fig 1B, C). Square-wave gratings at 12 directions were presented in pseudo-random order for 5 seconds each, interleaved with 3 seconds of mean-luminance matched grey screen. We designed grating stimuli with slightly longer durations than many studies [16, 29–30] for two reasons: first, to allow for the slow decay of the calcium indicator to fall to baseline in order to remove any confounds from the previous grating, and second, to study the sustained response in the population, rather than a transient response to stimulus onset [31]. Mice were awake and allowed to freely run on a linear treadmill. The majority of neurons showed significantly increased activity to one or more gratings over grey screen (3023/4535). Of the responsive subpopulation, most neurons were significantly tuned to orientation or direction (2073/3023; 540/3023 respectively). Neuron tuning was measured by fitting an asymmetric circular Gaussian tuning curve to the trial-averaged mean fluorescence in each grating direction (Fig 1D). These numbers of tuned and untuned neurons are in line with other population studies of awake, mouse V1 [16, 30]. In subsequent analyses, we pooled all visually responsive neurons without significant direction or orientation tuning into a class of ‘untuned’ neurons, differentiating two distinct subpopulations in V1 by their responsivity to drifting gratings.

**Fig 1.**
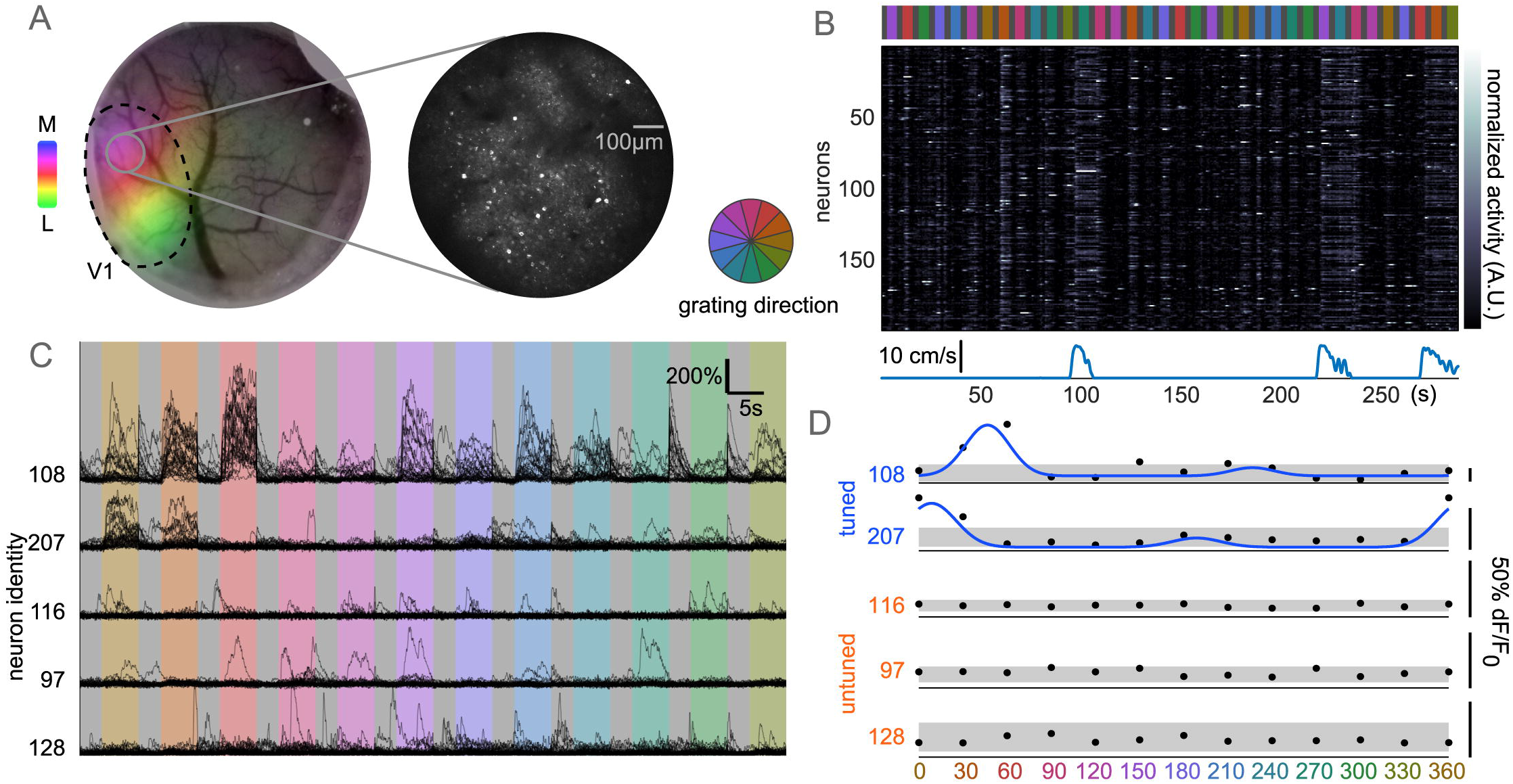
Categorizing response properties in V1 populations (A) Left, anecdotal cranial window, V1 retinotopy mapped by intrinsic signal overlaid (colormap shows visual field azimuth, lateral to medial). Right, two-photon field of view L2/3 population from grey region on left. (B) One block of stimulus presentation (top; grey interleaved gratings) with simultaneous imaging data showing neuron normalized fluorescence over time (middle). Bouts of running (bottom) and gratings show coordinated increased activity across neurons. (C) Five example neurons’ activity, black traces represent one trial, organized by grating direction. Grey period preceding gratings are kept together, with discontinuities at end of gratings. Top two neurons show increased activity to specific directions, others are untuned with substantial trial-to-trial variability. (D) Trial-averaged activity reveals classic tuning properties in a subpopulation of neurons. Tuned neurons fit with asymmetric circular Gaussian tuning curve (blue), untuned neurons show no increased activity relative to grey periods (grey bar mean+/-std average response).

### Responses are highly variable across trials

Single trial responses to gratings showed a high degree of variability, even in strongly tuned neurons, manifesting as occasional strong responses to null-directions and weak or absent responses to preferred directions (Fig 1C, 2A). On average, tuning curves described average response strength of tuned neurons well (0.70+/−0.20 R^2^). Responses across single trials in tuned neurons, however, were not well-described by their tuning curves. The mean fluorescence for each direction only explained a small fraction of the total trial-to-trial variance (Fig 2B). This finding matches earlier results in awake, mouse V1 [30]. We computed the distribution of response strength (mean fluorescence across a grey or grating presentation) within single trials, z-scoring to account for neurons with different activity levels. Single trial response distributions were skewed, with most responses weaker than the mean (Fig 2C). Tuned neurons showed slightly stronger responses during gratings, as compared to untuned neurons which had nearly identical response distributions in grey and grating trials. Tuned response distributions were strongly overlapping, however, consistent with the hypothesis that individual neuron activity is not solely driven by tuning properties.

**Fig 2.**
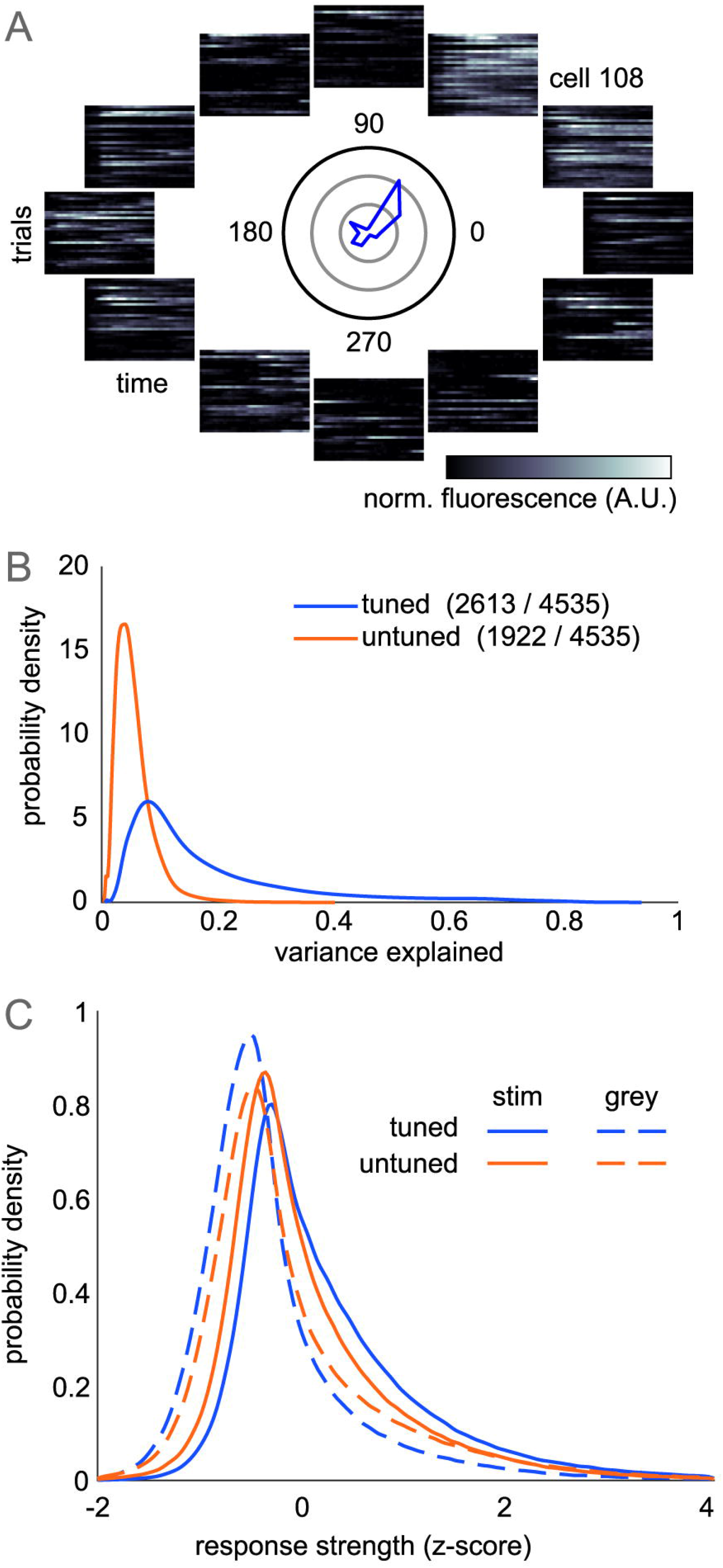
Single trial response variability (A) Single trial responses of representative tuned neuron. Mean tuning curve in polar coordinates shown in center (rings=100% dF/F_0_). Each heatmap positioned by grating direction shows normalized time-varying activity for each trial. (B) Trial-averaged tuning responses do not explain trial-to-trial variability. Tuning average (blue curve in A) is subtracted from mean fluorescence in each trial to compute variance explained across trials. (C) Averaging across time, neurons show highly variable responses during grating and grey trials with a right-skewed distribution due to strong activity in a few trials. Tuned neurons show larger increase in response during gratings than untuned neurons.

To further describe population activity, we computed the time-varying activity during the presentation of a grating and its preceding and following grey presentations. For each trial, we removed neurons that were silent (defined as no fluorescence change 2*S.D. above baseline), and computed the time-varying z-scored fluorescence across neurons (Fig 3A). Despite the long duration of stimulus presentations, adaptation effects were minimal in these L2/3 neurons, as tuned neurons showed sustained activity throughout the 5 second stimulus presentation. Untuned neurons have weak modulation to the onset and offset of the stimulus, and are equally active during the grey period. However, these effects are small in comparison to variability across trials, as indicated by strong overlap between the activity of the two subpopulations. In the awake animal, running speed is known to strongly influence spike rates [32], and we similarly observe that periods of high population activity are very likely to occur during periods of running (Fig 3B, anecdotally in 1B). However, mice did not preferentially run during grating or grey presentations (probability of running 9.7+/−7.7% during gratings; 10.0+/−7.4% during greys; p=0.278 paired t-test). While we used changes in fluorescence for all other analysis, we expanded our comparison of both subpopulations by estimating spike rates using a spike-inference from calcium fluorescence algorithm [33]. We found that untuned neurons exhibited an identical firing rate distribution to tuned neurons. Therefore, differences between subpopulation dynamics cannot be explained by differences in firing-rates.

**Fig 3.**
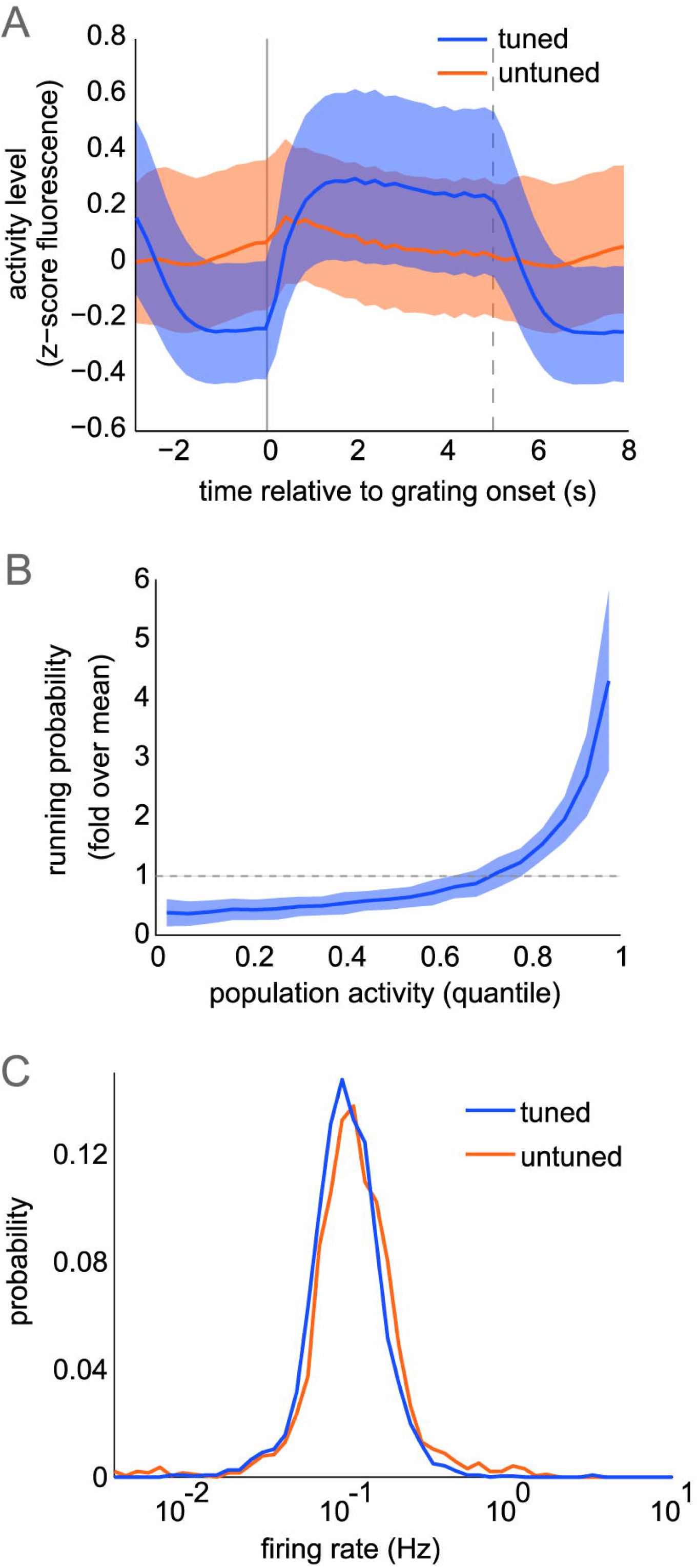
Population response properties (A) Mean time-varying fluorescence of active neurons, pooled across datasets, during grating stimulus and preceding, following grey periods in tuned, untuned subpopulations (median+/-quartiles across pooled neurons). Grating onset, offset shown in solid, dotted lines respectively. (B) Total population fluorescence split into equal quantiles to compute probability of running (speed>0.1cm/s). Plot shows mean+/-std across datasets (n=23). (C) Firing rate distributions computed across all imaging frames from spike trains inferred from fluorescence.

### Correlation structure during drifting grating and grey screen trials

Untuned neurons are a large proportion of total neurons, exhibit similar spike rates to tuned neurons, and are likely to contribute to correlation structure in the population. To begin describing how dynamics are affected by stimuli in populations containing tuned and untuned neurons, we first analyzed pairwise correlations within grating and grey presentations. We computed average correlation coefficient of the mean fluorescence within trials across neuron pairs without removing signal-dependent responses (i.e. signal correlations). Overall, within-subpopulation correlations are weak (0.014+/−0.027 tuned; 0.033+/0.047 untuned), and beween-subpopulation activity is slightly anti-correlated (-0.019+/−0.017). Comparing mean pairwise correlations between gratings and grey frames, only within-subpopulation correlations are affected by grating stimuli, while between-subpopulation correlations are constant between stimulus and grey conditions (Fig 4A). Tuned neurons show a strong decrease in mean correlations during gratings, as seen in macaques [34]. Conversely, untuned neurons are more strongly correlated during gratings, and yet correlations between tuned and untuned neurons do not change in magnitude between stimulus and grey conditions. Untuned neuron activity is not directly modulated by the stimulus, so changes within this subpopulation most likely reflect changes in activity from the tuned subpopulation propagating through local synaptic connectivity. However, this occurs without a change in mean correlation between tuned and untuned neurons, bringing to question the mechanism involved. The pairwise correlations in tuned neurons during the stimulus are a function of their preferred grating directions, as expected for signal correlations (Fig 4B). Similarly tuned neurons show strong correlations, while orthogonally tuned neurons show negative correlations. This structure is not present during activity in grey periods, however. This is surprising, because if local connectivity underlies correlations in the grey condition, one should expect the structure seen during gratings to remain in part because similarly tuned neurons are more likely to be connected [35].

**Fig 4.**
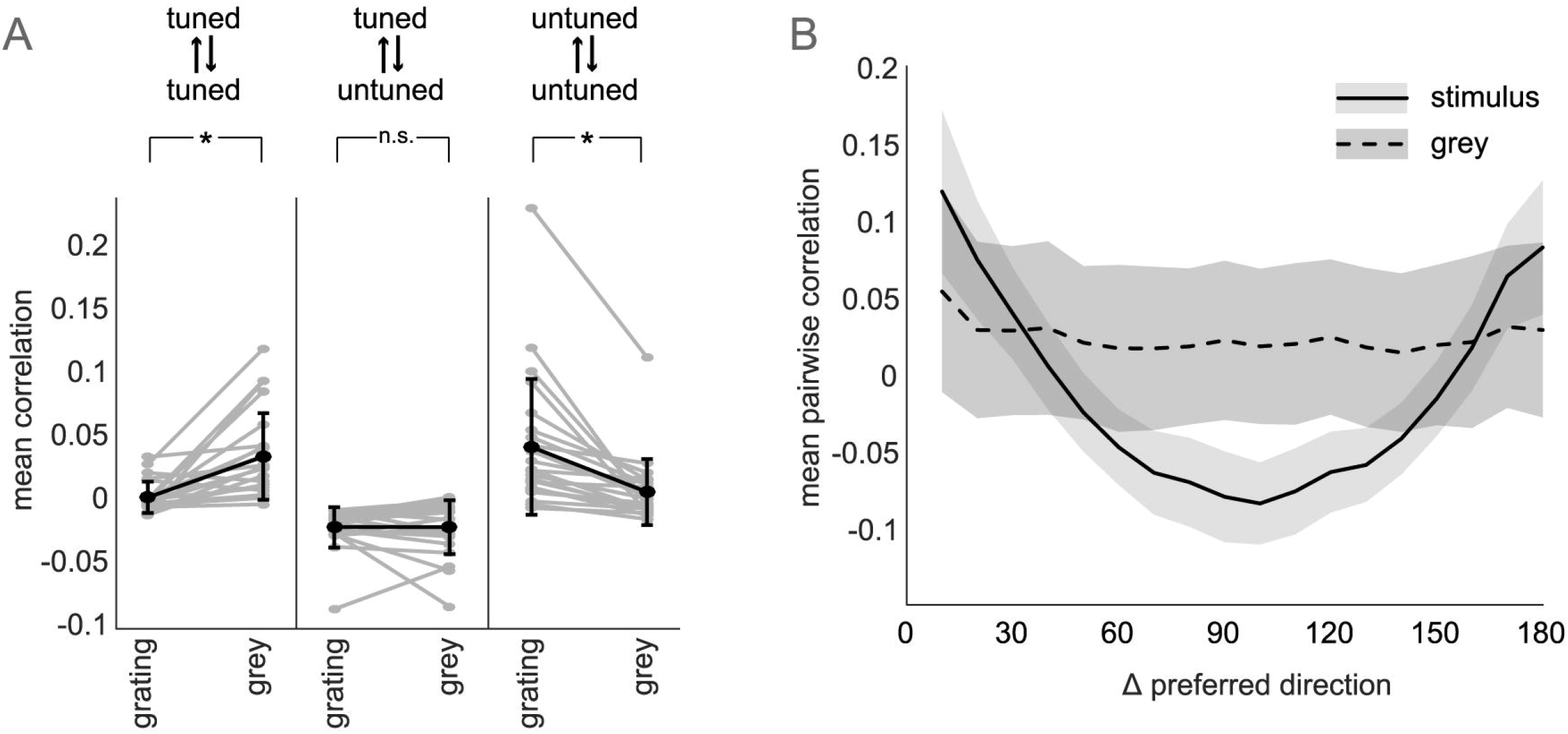
Correlations during grating and grey stimuli (A) Mean activity correlation between stimulus conditions, within and between subpopulations (mean+/−std in black; individual datasets in grey). Significance determined with Tukey-Kramer corrected one-way ANOVA at alpha=0.01 (tuned-tuned p=0.004; tuned-untuned p=0.999; untuned-untuned p=0.001) (B) Correlations within grating and grey trials for tuned neurons (mean+/−variance pooling neurons with given difference in tuning over datasets).

### Measuring trial-to-trial covariability

Overall correlations in populations including untuned neurons begins to reveal properties of local population activity, but in order to study the sources and structure of trial-to-trial shared variability, researchers attempt to remove the stimulus-dependent portion of responses leaving only variability, or ‘noise’ [20]. Correlated fluctuations between the remaining responses are therefore often called ‘noise correlations.’ In these data, however, the untuned subpopulation has no stimulus-dependent response, and the stimulus-driven response explains only a small portion of overall variability in tuned neurons (2B), making traditional noise-correlation analysis unsatisfactory. Furthermore, mouse V1 populations are characterized by global cofluctuations common to every neuron [22]. We therefore used the following partial-correlation analysis that accounts for stimulus-driven responses as well as population-wide cofluctuations in order to study pairwise noise correlations within and between subpopulations.

Gratings were presented in 5-minute blocks, each block contained three repetitions of each direction in pseudo-random order, and this order is maintained between blocks. We computed a partial correlation coefficient in each block between the activity of every pair of neurons, controlling for stimulus responses and population co-activity (Fig 5A). For each pair of neurons, the average activity across all remaining blocks represent the two stimulus-dependent responses capturing tuning properties, if present.

**Fig 5.**
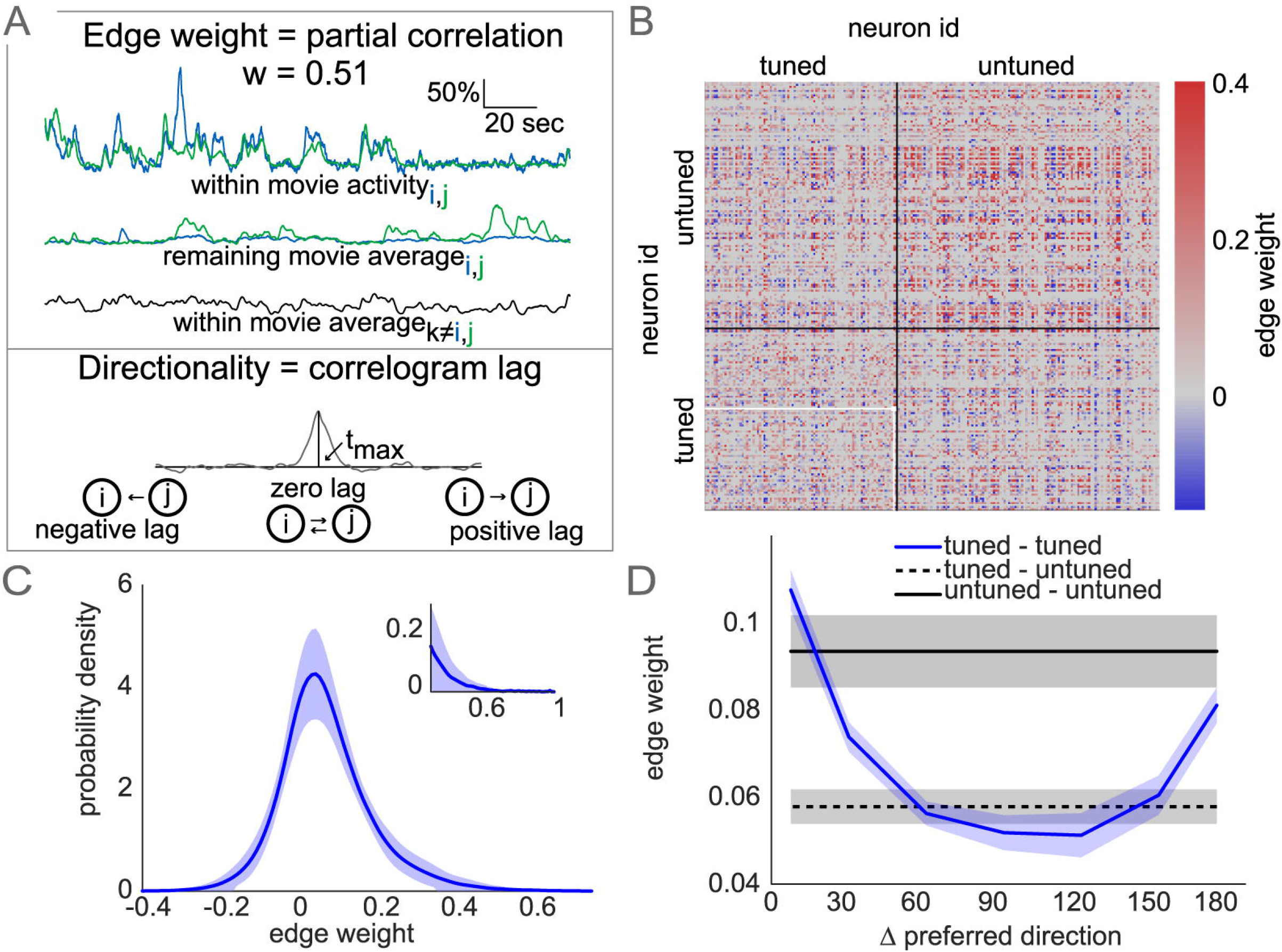
Partial correlation matrices representing trial-by-trial covariability (A) Edge weight estimated by mean partial correlation. Fluorescence traces of a neuron pair (i, j) within one movie, and three variables accounted for in partial correlation: remaining-movie average activity of i, j and within-movie average activity of remaining population (top). Edge direction set by offset of peak (t_max_) in neuron i, j correlogram (mean across movies). (B) Partial-correlation matrix sorted by subpopulation showing dense within- and between-population correlations. Neuron pair entry from (A) in white. (C) Kernel-density estimate distribution of all edge weights (mean+/−std across datasets). Inset shows right-tail of the strongest edges. (D) Tuned correlations match overall stimulus correlations, despite accounting for stimulus response, and slightly weaker between-subpopulation correlations (mean+/−sem over datasets).

The mean within-block population activity of all remaining neurons captures population-wide cofluctuations, for example running-speed effects. The mean partial-correlation across blocks is taken as the final correlation strength. Additionally, we added directionality to the partial-correlation by examining the mean cross-correlogram across blocks for the neuron pair. If the peak value occurs at lag 0 (i.e. within the same imaging frame), the edge was bidirectional, otherwise the edge was in the direction of positive lag. Lags greater than 500ms were thrown out and correlations set to zero. Though we interpret these partial-correlations as equivalent to noise correlations, the correlation matrices are different from traditional noise correlations in two important ways. First, many pairs of neuron correlations are exactly zero (51.7+/−7.2%), and second, non-zero correlations are asymmetric (Fig 5B). This allows us to analyze these matrices from a graph-theoretic perspective representing the matrix as a weighted, directed graph. Graph representations of pairwise edges allow us to analyze population-wide statistical features of the correlation structure. Overall, partial-correlation strengths were long-tailed, centered slightly above zero (Fig 5C), similar to noise-correlations observed elsewhere [36]. As expected, tuned noise correlations were similar to signal correlations [36–37], with similarly tuned neurons having stronger noise correlations on average (Fig 5D).

### Graph structure of partial-correlations

We next analyzed the partial-correlations within and between tuned and untuned subpopulations. The graphs exhibit dense correlations with varying strengths among subpopulations (Fig 6A). To analyze biases in edge strengths, we thresholded the matrices at increasing values, setting all edges below each threshold to zero. Among all edges, within-tuned connections are more likely, while within-untuned and between-subpopulations are slightly less common (Fig. 6B; within-tuned 54.2+/−8.6%, within-untuned 44.1+/−6.0% between 43.9+/−6.2%). Between-subpopulation connections remain the least likely at higher thresholds, but among the strongest edges, within-untuned connections are the most likely. We then recomputed partial-correlation matrices, exclusively using frames during the grey condition or during grating condition to see how correlation strengths were affected, despite controlling for mean stimulus-dependent activity. The two resulting matrices did not have significantly different edge strength (stimulus 0.12+/−.02; grey 0.12+/−.03; p=0.75 one-way ANOVA). However, probability of connection was higher within grey frames (grating 45.3+/−6.5%; grey 58.0+/−8.2%; p=5.8*10^-7 one-way ANOVA), indicating a higher degree of interconnectivity in the population in the absence of stimuli. When analyzing magnitude of edge strength differences between grey and grating matrices, we found that all connection types changed similarly with a mean near zero (Fig. 6C; within-tuned −0.011+/−0.097; within-untuned −0.005+/−0.103; between −0.008+/−0.093).

**Fig 6.**
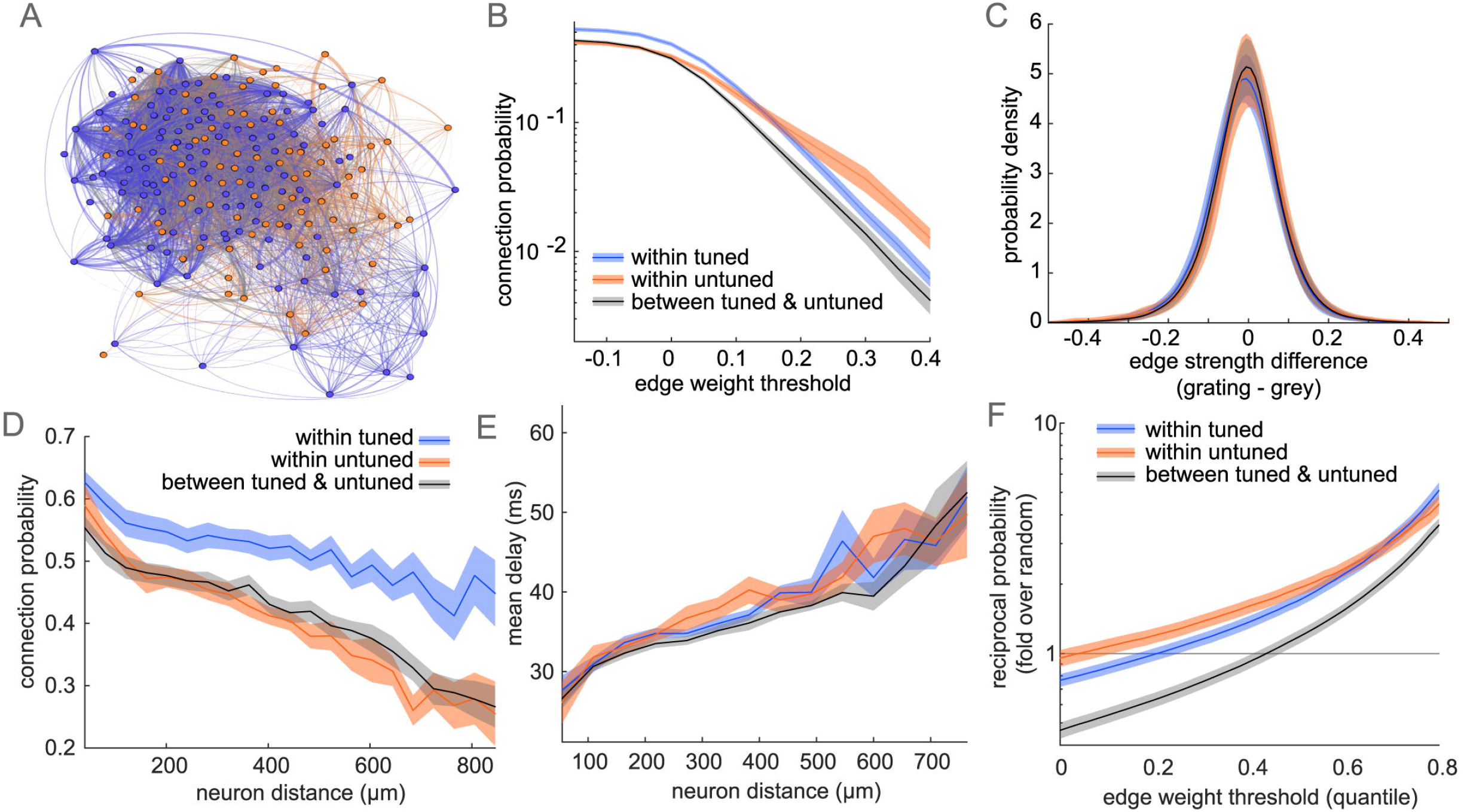
Directed graph structure of partial-correlation matrices (A) Example graph representation of matrix in (5B), nodes colored by subpopulation, edges colored by connection type as in (B-F), edge thickness represents absolute-value correlation strength. (B) Probability of connection by connection type for matrices with edges weaker than an increasing threshold set to zero. (C) Difference in edge strength between matrices using exclusively grating or grey frames. (D) Connection probability as a function of distance between neurons. (E) Mean lag of edges over neuron distance (F) Probability of a bidirectional edge at increasing thresholds, normalized by bidirectional probability assuming independence of edges. Plots in (B-F) show mean+/−sem across datasets.

Since population dynamics are partly constrained by synaptic connectivity [38], we evaluated whether there was a spatial component to the adjacency matrices. Edge probability fell monotonically with distance between neurons, similar to traditional noise correlations [36] and synaptic connections [8, 39]. Notably, the decay in connection probability is slower within tuned neurons compared to within untuned neurons (Fig 6D). Between-population decay lies in the middle. If the spatial structure of untuned correlations is exclusively driven by local connectivity, then bottom-up sensory drive is the most likely source for the longer range of functional correlations among tuned neurons. Furthermore, the mean lag (delay of the cross-correlogram peak used to determine edge directionality) is greater over longer distances and accumulated evenly across subpopulation connection type (Fig 6E). Assuming a linear change in lag over space, these data suggest the speed of functional correlations in this preparation is roughly 25 mm/s. As a function of edge strength, zero-lag edges (i.e. bidirectional edges) are more prominent within-subpopulations, and are strongly biased toward the strongest weights (Fig 6F). Thresholding at increasing edge strengths sparsifies the matrices, so we normalized the bidirectional edge counts by the probability of bidirectional edges assuming connections are placed randomly. Among all edges, bidirectional edges occur less often than random. The strongest edges, however, are roughly 5 times more likely to be bidirectional than random. Overall, zero-lag connections are less frequent between tuned and untuned neurons, suggesting a transmission or propagation of information, rather than simultaneous representation of the same information between subpopulations.

### Modeling trial-to-trial variability from local population activity

Because trial-averaged tuning poorly captures trial-to-trial responses we asked whether information in the local population could better explain V1 trial-to-trial response variability. Pairwise correlations have been shown to capture a significant portion of the complexity in population activity [17]. We tested whether the activity of a neuron could be modeled from the activity of its neighbors that had a non-zero correlation. Since correlation coefficients capture the linear relationship between neuron coactivity, we used a simple a linear combination of the input neuron activity with partial-correlation coefficients (edge strengths) as weights. This model gave a time-varying prediction of the fluorescence of a given neuron, which was then rescaled by an offset and a gain to account for different numbers of input neurons (Fig 7A). In many cases, this model resulted in highly accurate predictions of activity. Mean squared error of the reconstruction was small, and often near optimal compared to weights estimated by regression (Fig 7B). Tuned neurons were slightly better-modeled on average than untuned neurons (tuned MSE 0.014 median, 0.037 inter-quartile range; untuned MSE 0.017 median, 0.046 inter-quartile range), possibly because of the additional stimulus-dependent information captured by other tuned inputs. We selectively removed either tuned or untuned input neurons to evaluate the relative contributions to model predictions. Reconstruction error increased more when removing within-subpopulation inputs, but the effect is weak and distributions across neurons strongly overlap (Fig 7C). Correlations between tuned and untuned neurons contribute substantially to predicting their time-varying activity. We next asked whether our model based on local population activity could also predict trial-averaged tuning properties. For tuned neurons, we used the modeled fluorescence traces to recompute the mean fluorescence in each grating direction. The average responses in direction space were added together to obtain a mean tuning vector. The modeled fluorescence had very similar tuning properties to the data, as measured by the cosine similarity between model and data tuning vectors (Fig 7D). Tuned neurons with the best predictions also had the most similar tuning vectors. Local population activity therefore contains information to capture trial-to-trial variability as well as trial-average stimulus response properties.

**Fig 7.**
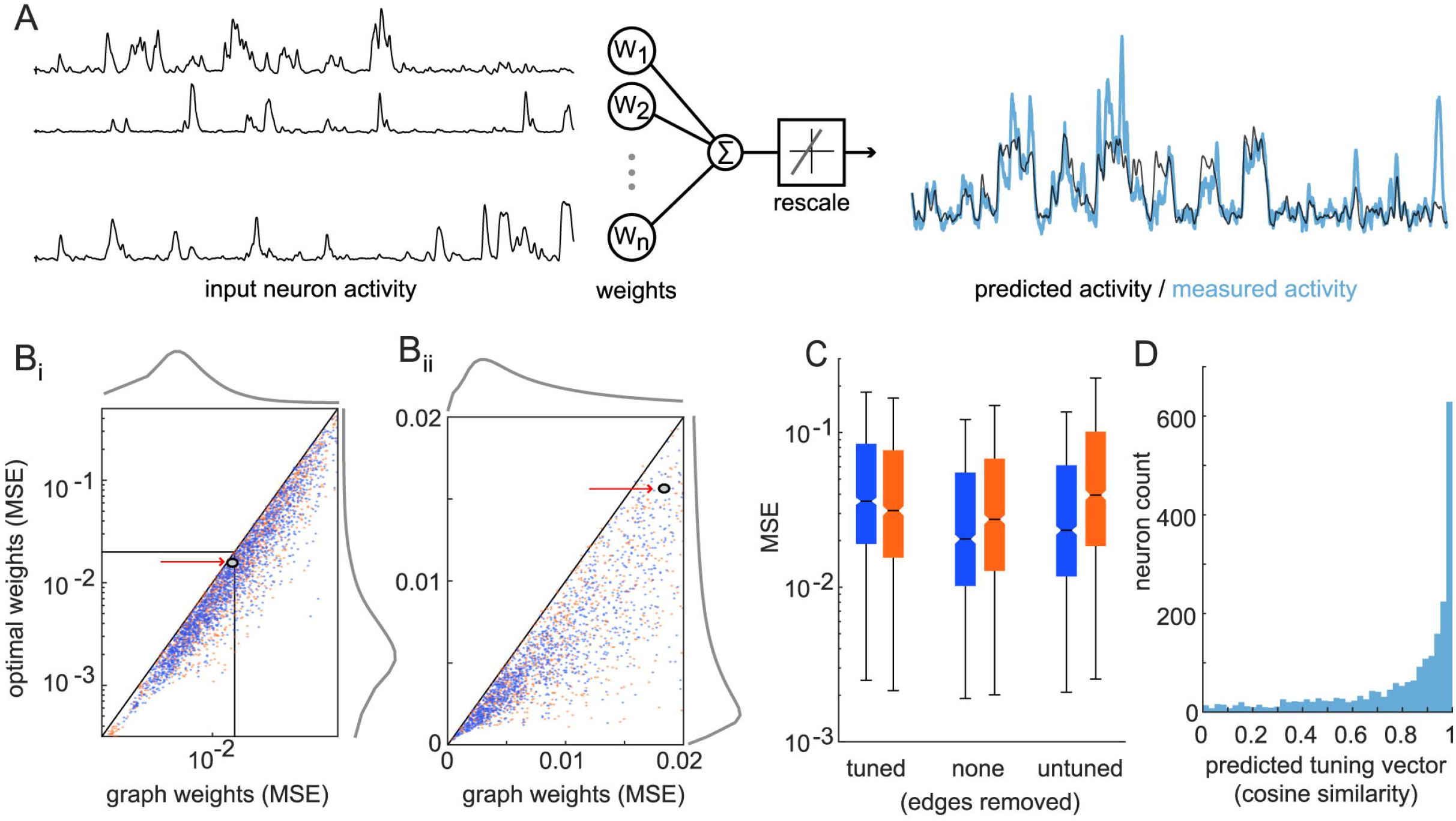
Modeling time-varying fluorescence from partial-correlation graphs (A) Illustration showing linear combination of incoming-edge neighbors fluorescence traces and rescaling to predict a given neuron’s activity. (Bi) Reconstruction quality across tuned and untuned neurons, quantified by mean-squared error (MSE) of reconstruction, compared to MSE using optimal weights (marginals in grey; arrows show neuron from (A)) (Bii) zoom in on neurons with best reconstruction. Lines show y=x solid; y=x/2 dashed; y=x/3 dotted. (C) Increase in MSE when selectively removing within- or between-subpopulation connections (boxplots show median, quartiles, with whiskers extending to most extreme value within 1.5*inter-quartile range outside quartiles). (D) Similarity of mean tuning vector between tuned neuron and reconstructed average activity.

### Pairwise and motif contributions to model performance

To investigate how individual neurons connections contribute to reconstruction of activity, we selectively removed input neurons based on their edge strength and measured the increase in reconstruction error (MSE), normalized by total mean squared fluorescence in the neuron. As expected, the strongest edges contribute the most to activity reconstruction with the strongest 25% of weights containing over half of the reconstruction capability, whereas removing half of the weakest edges had no discernible effect (Fig 8A). Interestingly, randomly removing edges maintains a nonlinear profile, suggesting possible synergistic effects between edges. Accounting for the cumulative weight removed, we still saw worse performance when removing the strongest edges first, and removing half of the total weight using only the weakest edges has minimal effect on reconstruction error (Fig 8B). Normalizing for the total weight removed reveals a nearly-linear increase in reconstruction error when removing the strongest weights, suggesting that these neurons may hold independent information from the remaining input pool.

**Fig 8.**
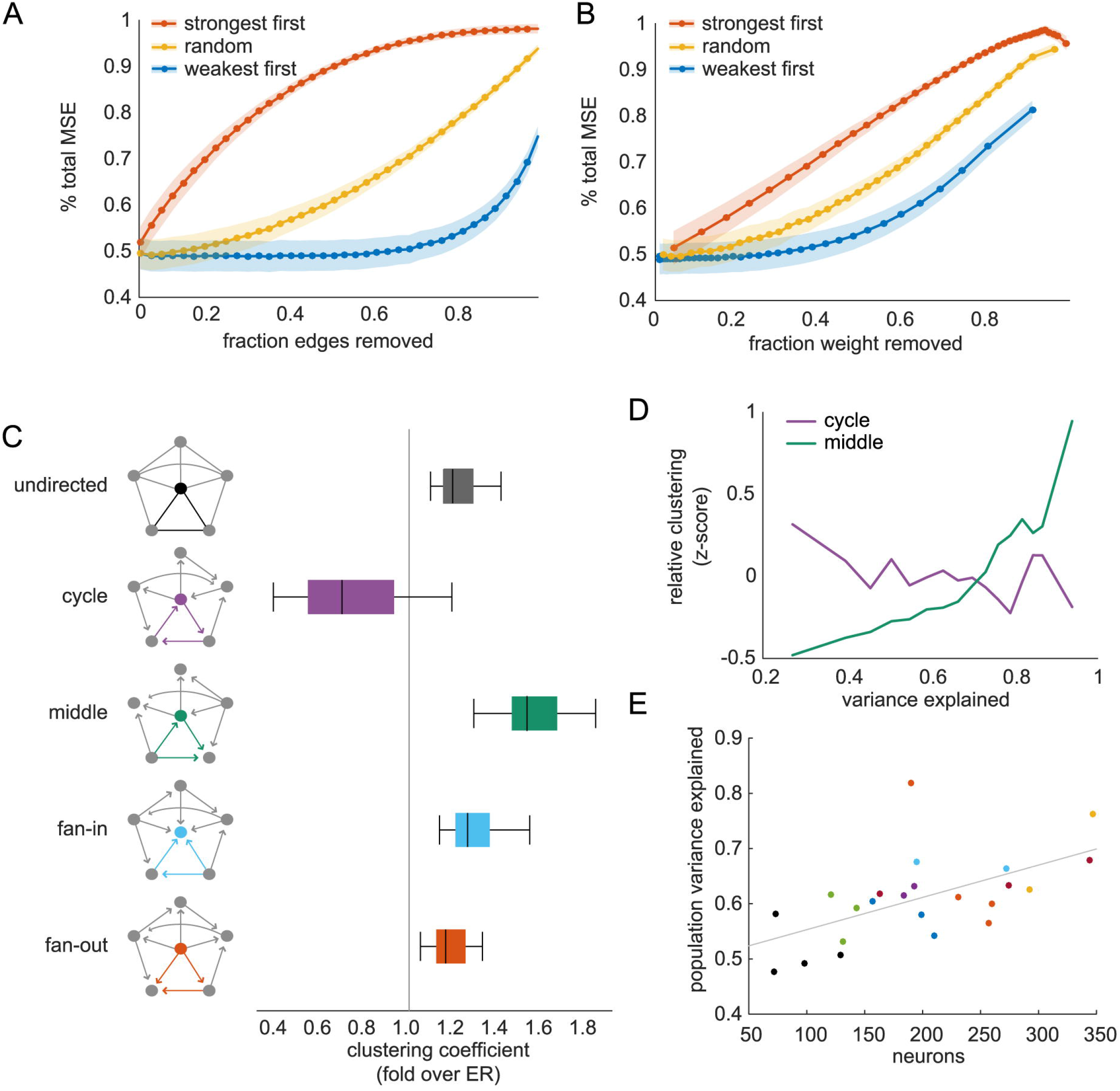
Pairwise and triplet motif contribution to model performance (A) Increase in mean-squared error (MSE) by number of edges removed, normalized by neuron total MSE (mean+/−variance across all neurons). (B) Increase in MSE by cumulative weight removed (mean+/−variance). (C) Clustering of triplet motifs in graphs (illustrated in left column) for each type of directed triplet motif. Box-plots as in Figure 7. Mean clustering was normalized by sparsity-matched Erdos-Reyni graphs. (D) Mean clustering coefficient, z-scored by mean/std for each motif type in (C), at different levels of model prediction showing neurons with best reconstruction had highest middle-man clustering. (E) Total population variance explained for each dataset (colored by mouse identity) shows better performance with more neurons imaged (linear trend in grey).

To address the possibility of synergistic or redundant information between input neurons, we analyzed connections between triplets of neurons termed ‘motifs’ in graph theory literature. Triplet connection motifs represent higher-order connectivity patterns that cannot be captured by individual edges, and can have strong implications for computation and information propagation within graphs [27]. We looked at the clustering of triplet motifs for each type of triangle that can be formed with directed edges (Fig 8C) [40]. Clustering is a measure of how many motifs are present among a neuron’s neighbors given its input and output connections. In comparison to Erdos-Reyni graphs, which have uniform, low levels of clustering across motifs, cycles of edges in the data are less clustered on average, with all other motifs showing elevated clustering. The middle-man motif shows the strongest clustering. Similar results have been found in activity generated by simulated and ex vivo neural networks, although fan-in clustering was higher than middle-man [23]. To address the possibility that triplet motifs are responsible for explaining more of the neuronal response than pairwise edges alone, we analyzed the relationship between motif clustering in single neurons and the variance explained by the predicted activity. Because the magnitude of clustering is different across motifs, we first z-scored clustering coefficients, then computed the mean across neurons with different levels of variance explained. Neurons with the best reconstruction showed higher clustering of middle-man motifs and lower cycle clustering, relative to other neurons in the population (Fig 8D). Total clustering, as well as fan-in and fan-out (not shown for clarity), had weak, positive correlation with variance explained. Together, these relationships map directly onto the overall prevalence of the graph motifs, suggesting that the graph structure has an important function in representing population information.

Because the motif with the highest mean clustering was also most indicative of model performance, we hypothesized that the partial-correlation structure might underlie our ability to predict neuron activity from its local population. Interestingly, across fields of view, the total variance explained in the population increased with number of neurons imaged (Fig 8E; r=0.58). This effect suggests that, in addition to motifs, total neurons sampled in the population determines our ability to measure a neurons’ single trial dependence on its local population. Moreover, the linear trend had no discernible plateau, so representing the local population may require recording from more than 300 neurons simultaneously.

## Discussion

### Summary and correspondence with previous findings

We sought to describe the interrelationships within local populations of V1 neurons, including tuned and untuned neurons, as they relate to single-trial responses to grating stimuli. We used two-photon imaging to record from L2/3 excitatory populations constitutively expressing calcium indicator GCaMP6s. Neurons with similar response properties showed stronger co-variability on average, but across the entire population there was a broad distribution of correlations driven by a confluence of sensory drive and activity in the local population. The functional correlations in the recorded populations were sufficient to predict activity in individual neurons, far surpassing predictions from tuning characteristics alone. The dependence on local population activity reinforces theories of layer 2/3 acting under strong modulation with sparse activity and weaker dependence on sensory drive than layer 4 [41]. We summarized the structure of correlations as directed, sparse matrices in order to analyze population dynamics from a graph theoretic perspective. We demonstrate that a simple population model capable of predicting single-trial neural responses is also able to accurately predict trial-averaged tuning responses, a key feature of V1 function. We found a specific triplet connectivity motif that correlated with our ability to predict activity on single-trials. This result could not have been observed from only studying pairwise correlations and motivates the continued use of graph theory to study neural population dynamics.

The single-neuron response properties in our data replicate imaging and electrophysiological results in awake recordings of V1 including proportion of tuned neurons [29-30, 36], and variance explained by the mean tuning curve [30]. Both tuned and untuned neurons have low firing rates on average, though estimation of firing rates from calcium imaging is not always straightforward. We note that single neuron properties, including firing rate and trial-to-trial variance, can change substantially between animal models and in different states of anesthesia [42–43].

### Population structure of pairwise correlations

We found that the magnitude of signal correlations between tuned and untuned subpopulations do not change between grey and grating stimulus conditions, while within subpopulation correlations do change. Perhaps more expected would be ubiquitous changes, or exclusive changes among neurons modulated by the stimulus. Within subpopulation correlations change in opposite directions, however, and could serve as a mechanism to balance changes between the subpopulations.

We found that the spatial organization of the network is a strong determinant of correlation structure with correlations decaying over distance, consistent with paired patch clamp recordings [8] and the correlation structure of activity in isolated preparations [44]. Correlation matrices were computed as an asymmetric partial correlation coefficient, accounting for stimulus and population effects, to allow the incorporation of untuned neurons into traditional noise correlation analyses. While this approach differs from standard noise correlation estimates, the magnitude of correlations and dependence on tuning similarity replicate previous results employing noise correlations [36]. For this reason, we interpret the partial correlations as measuring trial-to-trial covariability, equivalent to noise correlations. Tuned neurons, which combine feedforward sensory inputs with recurrent inputs, show a slower spatial decay of correlational values than untuned neurons, the latter of which presumably are driven more exclusively by recurrent, local inputs. These data show than subdividing the population by their response to grating stimuli can differentiate rates of spatial decay within the network.

Noise correlations are hypothesized to be driven in part by shared synaptic input [45]. Spatial dependencies of feedforward and recurrent connectivity can qualitatively change the spiking activity in network models [46]. The relatively small diameter of the fields of view imaged here (<1mm vs 10mm) does not allow us to differentiate between the two modes of activity predicted by these models. These data suggest, however, that functional recurrent correlations have less spatial extent than feed-forward functional correlations. To our knowledge, analogous synaptic connectivity estimates do not exist for mouse V1.

In addition to spatial structure of correlations, we found an increase in mean temporal delay of correlations over distance. This delay spans roughly 20-50 ms in our field of view and is therefore likely to reflect timescales of functional correlations rather than monosynaptic transmission delays. The implied speed of this delay accumulation is much faster than propagating LFP waves observed in macaque M1 [47] after normalizing for total cortical surface area [48]. The propagation of beta-oscillations and temporal delays of functional correlations likely have different underlying mechanisms, which may explain the different speeds.

### Prediction of single-trial responses from pairwise correlations

We were able to extract a substantial amount of predictive information from the local population using a simple, linear model. Alternative models incorporating known neuronal nonlinearities [49], or more sophisticated or biologically-relevant predictive models [50], may have improved performance. Consequently, the increase in total variance explained with population size may show different trends with alternative models. Nevertheless, we chose this modeling approach to maintain ease of interpretation and utilize the linear correlation coefficients estimated from the data in a straightforward manner. The linear models had good performance, and allowed us to remove input neurons without laboriously re-training the models. This straightforward modeling approach may not capture all of the information present in the local population, but its performance sets a lower bound on predictive information in the population. A small subset of neurons have high mean-squared error of predictions, and we speculate that we have not sampled enough of the population to predict these neurons given that total variance explained scales with population size. Neuron activity was only predicted from the in-degree neurons, rather than the entire population, reducing the number of parameters by 52+/−22% across neurons, though in either case, parameters scale by order N. These results substantiate previous models showing that the collection of pairwise relationships in large populations can explain complex activity patterns [17]. We further explored this idea by identifying a specific motif of pairwise correlations underlying accurate predictions of neuron activity on single-trials.

### Functional consequences of triplet motifs

The finding that middle-man motifs underlie the best predictions of activity may be initially surprising from the connection pattern of the motif. Compared to the fan-in motif, which has two input connections, the middle-man has one input and one output connection, and acts to route connections from its input to its target. However, the motifs were quantified using the clustering coefficient, which normalizes motif count by the total number of possible motifs (i.e. high fan-in clustering doesn’t correspond to high in-degree). The functional significance of any given triangle motif clustering is unknown and is likely dependent on the underlying system represented by the graph. In a neural network, middle-man clustering may indicate a hub-like property common in the brain [51], having a combination of convergent and divergent correlations. The cycle motif similarly has a combination of convergence and divergence, yet its clustering has a negative effect on prediction accuracy. The difference in the two motifs is the direction of connections between neighbors. In these networks, it may be that cycle clustering reflects recurrent, redundant correlations reducing our ability to predict activity from the population. Conversely, the middle-man motif is isometric to fan-in and fan-out motifs, and could allow for transfer of information between motifs, increasing predictive power. Finally, this result demonstrates that in addition to providing insights into synaptic mechanisms underlying dynamics [23], network science can also provide insights into predictions of single trial neural responses as we have demonstrated here.

V1 is the first stage in which visual information is encoded in densely recurrent cortical networks. Thus, in order to understand activity patterns in V1, one must take into account visual drive as well as local network activity. We have provided a quantitative comparison of the relative influences of these two factors in awake, ambulating mice. Local network effects dominate on single trials, highlighting the importance of investigating cortical computation from a population perspective in order to understand how information is encoded in single trials. Populations of neurons exhibit emergent properties beyond the sum of their individual neurons and connections, and we use the analytic framework of graph theory to begin unraveling this emergent structure.

## Materials and Methods

### Animals and surgery

All procedures were performed in accordance and approved by the Institutional Animal Care and Use Committee at the University of Chicago. Data was collected from C57BL/6J mice of either sex (n=4 female; 4 male) expressing transgene Tg(Thy1-GCaMP6s)GP4.12Dkim (Jackson Laboratory) between ages P84 – P191. After induction of anesthesia with isoflurane (induction at 4%, maintenance at 1-1.5%), a 3mm diameter cranial window was implanted above V1 by stereotaxic coordinates and cemented in place alongside a custom titanium headbar. Mice recovered for at least 8 days before intrinsic signal imaging to identify V1 followed by two-photon data collection.

### Intrinsic Signal V1 identification

Boundaries of V1 were identified by intrinsic signal imaging post-surgery [52] (Fig 1A, left]. Mice were anesthetized with isoflurane and head-clamped under a CCD camera (Qimaging Retiga-SRV). A vertical white-bar stimulus (100% contrast, 0.125Hz) was repeatedly presented on an LED monitor (AOC G2460) approximately 20cm from the contralateral eye while capturing cortical reflectance under 625nm illumination. The retinotopic mapping of V1 was then estimated at each pixel from the phase of peak reflectance driven by increases in activity-dependent blood flow.

### Data collection

Two-photon imaging was collected from awake, head-fixed mice on a linear treadmill. Running speed was measured with a rotary encoder attached to the treadmill axle. A L2/3 field of view (roughly 800μm diameter) in V1 was identified with galvanometer-mirror raster scanning (Cambridge Technologies; 6215H). Once a suitable field of view was found, raster scans (1Hz) were continuously acquired for roughly 10 minutes alongside visual stimulation. Neurons (n=72–347 per field of view) were then automatically identified using custom image processing software for imaging during visual stimulation using Heuristically Optimal Path Scanning [53] at 25–33 Hz). All imaging was performed at 910nm (Coherent; Chameleon Ultra) with a 20X 1.1NA Olympus objective and GaAsP PMT (Hamamatsu; H10770A-40). Field of view size was estimated by fitting circles to a single raster scan of 15μm fluorescent microbeads and used for each dataset, though true field of view size may vary up to 8% across datasets from realignment of laser beam path.

### Visual presentation

Drifting grating stimuli were presented on an ASUS VG248QE, 20cm from the contralateral eye at 60Hz; 60cd/m2. The mean luminance was measured and gamma correction was performed and confirmed using a luminance meter. Square-wave gratings were shown at 80% contrast, 2Hz, 0.04 cyc/deg at 12 evenly spaced directions. Gratings were presented for 5s, interleaved with 3s mean-luminance grey screen. Three repetitions of each orientation were presented in a pseudo-random order, resulting in a roughly 5min stimulus movie. The grating order was preserved between movie presentations, and mice were shown 8-11 repetitions of the movie (24-33 repetitions of each direction).

### Data acquisition and pre-processing

Stimulus presentation was monitored with a photodiode (Thorlabs) and synchronized with running speed and imaging frames at 2kHz. For each neuron, baseline fluorescence was estimated from raw fluorescence by thresholding to eliminate spike-induced fluorescence transients and smoothed with a 4th-order, 81-point Savitzky-Golay filter. Fluorescence time-series were then normalized to percent change from baseline (dF/F_0_) using this time-varying baseline. Spike inference from fluorescence traces was performed using the OASIS algorithm [33] implemented using software made freely available (github.com/j-friedrich/OASIS). Inference outputs probability of spiking at each time point. As commonly done [54], probabilities were thresholded to obtain a binary spike train.

### Response properties and tuning classification

Neurons were classified as visually responsive if the mean response to any grating was significantly greater than the mean response across all grey periods by Dunnett-corrected one-way ANOVA (alpha = 0.01). In these analyses, each trial response is the mean fluorescence across the entire grating presentation, or the last half of the grey presentation to allow for fluorescence from grating responses to decay. Responsive neurons were then tested for statistically significant orientation- or direction-tuned responses according to the trial vectors in orientation or direction spaces ([55] for more detailed methods]. For significantly tuned responses, tuning curves were then fit with an asymmetric-circular Gaussian to significantly tuned neurons. Tuning curve parameters (baseline, tuning width, peak amplitudes, and preferred direction) were fit repeatedly using randomized initial conditions. The parameter set that minimized mean-squared error was maintained.

### Signal correlations

For each neuron, the mean responses in each trial (using the same time windows as for tuning classification) for either stimulus or grey trials were used as response vectors. The correlation coefficient between each pair of these vectors was used to compute a correlation matrix for the grating and grey conditions.

### Partial correlation matrices

For each pair of neurons, pairwise-correlation was computed as the mean partial correlation between their fluorescence across movies while accounting for three variables. For a matrix with all five variables as columns, the two neuron fluorescences, *x*, and the three control variables, *z*, the covariance matrix S is

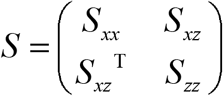

Then the partial correlation coefficient is normalized from ρ with

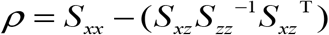

Here, a movie is the 5-minute block of grating presentations that was repeated in each experiment. The first two variables are the mean response of the two neurons in all other movies, accounting for the stimulus-driven responses. This is equivalent to normalizing by the mean response as in traditional noise correlation estimates. The within-movie mean fluorescence of all other neurons was also included to control for population-wide covariability, for example running speed effects. Furthermore, the mean cross-correlogram across movies was used to compute directionality of the correlation. The time-lag of the cross-correlogram global maximum determined the direction and lag of the edge. If the lag was zero, the correlation was bidirectional. If the lag was greater than 500ms (roughly 14 imaging frames), no edge was included.

### Graph analysis

The partial-correlation matrix could equivalently be analyzed as a directed, weighted graph. Open source software (Gephi) was used for visualization, with node layout determined by the Yifan-Hu algorithm and tuned by hand. Edge weights less than 0.05 were set to zero for visualization clarity. Erdos-Reyni (ER) null graphs were generated for each graph to match the mean connection probability. The mean directed clustering coefficient across nodes was calculated across 50 ER graphs and averaged for comparison with data. Clustering coefficients were computed with binary matrices (nonzero weights set to one).

### Modeling neuron fluorescence

To model neuron responses, we used a linear weighting of the fluorescence of every in-degree using the weights in the partial-correlation matrix. At each time point, a weighted sum was calculated, resulting in a time-varying predicted fluorescence trace. Because different numbers of input neurons and varying calcium transient amplitudes, the modeled trace was then fit with a linear offset and a gain to minimize mean-squared error with the true fluorescence trace. These two parameters were not changed when input neurons were removed (7C, 8A, 8B). For tuned neurons, we also recomputed the trial-averaged tuning response of modeled activity. The fluorescence over a grating presentation was averaged, then trials were averaged over directions to obtain a mean direction-response. The sum of these vectors in direction space gave the model-estimated tuning vector, and the cosine similarity with the data-derived tuning vector was used to quantify the reconstruction of the tuning properties. Cosine similarity was computed in direction or orientation space according to each neuron’s tuning properties. To compute total population variance explained, modeled traces were subtracted from the data traces to obtain residuals, and the ratio of variances were subtracted from one as

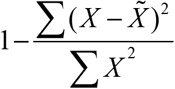

### Optimal weight estimation

Optimal weights for all incoming edges were computed for each neuron by LASSO regression as

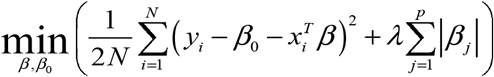

For weights β, and offset β_0_, with modeled neuron fluorescence as y and input neuron activity as x. Weight estimation and 5-fold cross validation to estimate MSE standard error was performed with MATLAB R2016a implementation. The maximum regularization parameter (λ) whose mean-squared error did not exceed the standard error of the minimum MSE was used to find the set of optimal weights.

## Acknowledgments

This work was supported by Big Ideas Generator (BIG) Vision Fund- From neural circuits to computation. We thank John Maunsell and Jackson Cone for critical feedback and discussion. We thank members of the MacLean lab for helpful comments on the manuscript.

